# Mice with Induced Pulmonary Comorbidities Display Severe Lung Inflammation and Mortality following Exposure to SARS-CoV-2

**DOI:** 10.1101/2020.10.28.358614

**Authors:** Reut Falach, Liat Bar-On, Shlomi Lazar, Tamar Kadar, Ohad Mazor, Moshe Aftalion, David Gur, Ohad Shifman, Ofir Israeli, Inbar Cohen-Gihon, Galia Zaide, Hila Gutman, Yentl Evgy, Yaron Vagima, Efi Makdasi, Dana Stein, Ronit Rosenfeld, Ron Alcalay, Eran Zahavy, Haim Levy, Itai Glinert, Amir Ben-Shmuel, Tomer Israely, Sharon Melamed, Boaz Politi, Hagit Achdout, Shmuel Yitzhaki, Chanoch Kronman, Tamar Sabo

## Abstract

Severe manifestations of COVID-19 are mostly restricted to people with comorbidities. Here we report that induced mild pulmonary morbidities render SARS-CoV-2-refractive CD-1 mice to be susceptible to this virus. Specifically, SARS-CoV-2 infection after application of low-doses of the acute-lung-injury stimulants bleomycin or ricin caused a severe disease in CD-1 mice, manifested by sustained body weight loss and mortality rates of >50%. Further studies revealed markedly higher levels of viral RNA in the lungs, heart and serum of low-dose-ricin pretreated, as compared to non-pretreated mice. Notably, the deleterious effects of SARS-CoV-2 infection were effectively alleviated by passive transfer of polyclonal or monoclonal antibodies generated against SARS-CoV-2 RBD. Thus, viral cell entry in the sensitized mice seems to involve viral RBD binding, albeit by a mechanism other than the canonical ACE2-mediated uptake route. In summary, we present a novel mice-based animal model for the study of comorbidity-dependent severe COVID-19.

## Introduction

Severe manifestations of COVID-19 are mostly restricted to distinct groups of people; elderly persons and such that have comorbidities, form a significantly high proportion of those which develop acute lung injury and acute respiratory distress syndrome (ALI/ARDS) and thereby require intensive care (*1, 2*). These observations suggest that the severe form of COVID-19 is an outcome of the combined effects of both SARS-CoV-2 infection and an underlying pathological condition or predisposition. In the laboratory most COVID-19 animal models, based on ferrets, hamsters and non-human primates, develop a mild pathology which resolves within a relatively short period of time (*3-5*) and thus seem to reflect the more prevalent asymptomatic-to-mild manifestation of the disease observed in humans. As such, these animal models do not enable a robust assessment of the therapeutic potential of candidate drugs and therapeutic antibodies which may alleviate the life-endangering form of the COVID-19 disease. Lately, to overcome the inability of SARS-CoV-2 to bind to murine ACE2, mice animal models in which human ACE2 is transiently or constitutively expressed were established (*6-8*). Although severe clinical manifestations were observed in some of these genetically-modified mice models (*8*), these were not dependent on pre-existing morbidities, as in the case of severe COVID-19 in humans. Establishment of a COVID-19 animal model in which severe clinical signs are manifested only in the presence of a pathological background may prove to be more translationally-relevant for the study of this disease.

Bacterial lipopolysaccharide and bleomycin are well-established stimulants of ALI/ARDS in animal models (*9*). Ricin, a plant-derived toxin, has also been proven in recent years to cause ALI/ARDS (*10-11*). We therefore examined whether pulmonary exposure of mice to carefully chosen sub-lethal doses of these stimulants could cause a relatively mild and transient lung injury which may serve as an underlying pathology that promotes increased susceptibility towards the SARS-CoV-2 virus.

In the present study we show that following infection with SARS-CoV-2 *via* intranasal application, low-dose ricin (LDR)- and bleomycin-pretreated mice display a significant and sustained decrease in body weight and that a significant proportion of these mice die within a surveillance period of 2 weeks. Passive transfer of polyclonal or monoclonal antibodies directed against the SARS-CoV-2 virus or the RBD portion of the viral spike resulted in full protection of LDR mice following viral infection. Our findings demonstrate that mice with induced pulmonary injuries can serve as an accessible model for studying SARS-CoV-2 pathogenesis and treatment, and imply a non-canonical mode of viral entry in comorbidity-dependent severe COVID-19.

## Materials and Methods

### Cells and virus

African green monkey kidney clone E6 cells (Vero E6, ATCC® CRL-1586™) were grown in Dulbecco’s Eagle’s medium (DMEM) containing 10% fetal bovine serum (FBS), MEM non-essential amino acids (NEAA), 2 mM L-Glutamine, 100 units/ml penicillin, 0.1 mg/ml streptomycin, 12.5 units/ml nystatin (P/S/N) (Biological Industries, Israel). Cells were cultured at 37°C, 5% CO_2_ at 95% air atmosphere.

SARS-CoV-2 (GISAID accession EPI_ISL_406862) was kindly provided by Bundeswher Institute of Microbiology, Munich, Germany. Virus stocks were propagated (4 passages) and titered on Vero E6 cells. Handling and working with SARS-CoV-2 virus were conducted in a BSL3 facility in accordance with the biosafety guidelines of the Israel Institute for Biological Research (IIBR).

### Ricin, bleomycin, LPS

Crude ricin was prepared by us, from seeds of endemic *Ricinus communis*, essentially as described before (*13*). Bleomycin sulfate, (from *Streptomyces verticillus*, Cat. No. B8416-15U) and lipopolysaccharide (LPS, from *Escherichia coli*, Cat. No. L4391) were purchased from (Sigma, Israel).

### Animal experiments

Animals in this study were female CD-1 mice (Charles River Laboratories Ltd., Margate, UK) weighing 27-32 grams. Prior to treatment or infection, animals were habituated to the experimental animal unit for at least five days. Treatment of animals was in accordance with regulations outlined in the USDA Animal Welfare Act and the conditions specified in the National Institute of Health Guide for Care and Use of Laboratory Animals. All mice were housed in filter-top cages in an environmentally controlled room and maintained at 21±2°C and 55±10% humidity. Lighting was set to mimic a 12/12 h dawn to dusk cycle. Animals had access to food and water *ad libitum*. Ricin (1.7 μg/kg), bleomycin (2U/kg) or LPS (1.7 mg/kg) were applied intranasally (25μl per nostril) to mice anesthetized by an intraperitoneal (i.p.) injection of ketamine (1.9 mg/mouse) and xylazine (0.19 mg/mouse). Mice displaying weight loss of >30%, were euthanized by cervical dislocation. SARS-CoV-2 diluted in PBS supplemented with 2% FBS was intranasally instilled to mice anesthetized as above. Where indicated, mice were administered (intraperitoneal injection, 1 ml) one of the following antibodies: equine-derived polyclonal anti-ricin F(ab)_2_ fragment (1730 IsNU) (*14*), monoclonal anti-SARS-CoV-2 RBD (MD65) antibody (1 mg) (*18*), polyclonal anti-SARS-CoV-2-RBD antibody (SARS-CoV-2 NT_50_= 1:20,000) or polyclonal anti-SARS-CoV-2 (SARS-CoV-2 NT_50_= 1:10,000) antibody. For generation of RBD-huFc-and monomeric RBD were expressed and purified as described (*18*). Polyclonal anti-SARS-CoV-2-RBD antibody was generated by immunizing a rabbit (female New-Zealand White) with RBD-huFc (150 μg, Complete Freund’s adjuvant) followed by boosting with RBD-huFc (150 μg, Incomplete Freund’s adjuvant) at day 21 and with monomeric RBD (150 μg, Incomplete Freund’s adjuvant) at day 42. Hyperimmune serum was collected at day 50. Polyclonal anti-SARS-CoV-2 antibody was generated in a rabbit (female New-Zealand White) by repeated intravenous injection of live SARS-CoV-2 virus (1×10^6^ PFU, at days 0, 7, 10, 14 and 17). Serum was collected 14 days after the last virus injection.

### Mice activity measurement

Nocturnal activity of mice groups was monitored, utilizing a computerized home cage monitoring system (HCMS100) with a single laser beam and detector as described previously (*13*).

### Clinical laboratory analysis

Differential blood counts were determined in peripheral blood. Samples were collected from the tail vein of mice into EDTA containing tubes (BD, Franklin Lakes, NJ, USA) and were analyzed using Veterinary Multi-species Hematology System Hemavet 850 (Drew Scientific, Miami Lakes, FL, USA).

### Bronchoalveolar fluid (BALF) analysis

BALF, collected by instillation of 1 ml PBS at room temperature *via* a tracheal cannula, was centrifuged at 1500 rpm at 4°C for 10 min. Supernatants were collected and stored at -20°C until further use. BALF levels of IL-6, IL-1β and TNF-α, were determined using ELISA kits, following the manufacturer’s instructions (R&D Systems, USA). Protein concentrations in BALF samples were determined by Nanodrop (Thermo scientific 2000 spectrophotometer, Thermo Fisher, MA, USA).

### Measurement of ricin catalytic activity

Ricin-induced depurinated 28s rRNA in mice lungs was measured as described previously (*28*) and expressed as percent of total 28S rRNA.

### Measurement of viral RNA

Mice lungs, trachea, nasal turbinate, heart, spleen, liver, kidney and brain were harvested and grinded. Serum was separated from collected blood. From each sample, 200 μl were added to LBF lysis buffer (supplied with the kit) and viral RNA was extracted using RNAdvance Viral Kit (Beckman Coulter) and further processed on the Biomek i7 Automated Workstation (Beckman Coulter), according to the manufacturer’s protocol. Each sample was eluted in 50 μl of RNase-free water. Real-time RT-PCR assays, targeting the SARS-CoV-2 E gene, were performed using the SensiFAST Probe Lo-ROX One-Step kit (Bioline). The final concentration of primers was 600nM and the probe concentration was 300nM. Primers and probe for the E gene assay were taken from the Berlin protocol published in the WHO recommendation for the detection of SARS-CoV-2. The primers and probe sequences were as follows:

E_Sarbeco_F1 ACAGGTACGTTAATAGTTAATAGCGT,

E_Sarbeco_R2 ATATTGCAGCAGTACGCACACA,

E_Sarbeco_P1 ACACTAGCCATCCTTACTGCGCTTCG.

Thermal cycling was performed at 48^0^C for 20 min for reverse transcription, followed by 95°C for 2 min, and then 45 cycles of 94°C for 15 s, 60°C for 35 s. Cycle Threshold (Ct) values were converted to calculated PFUs with the aid of a calibration curve tested in parallel.

### Viral growth kinetics

Viral growth kinetics were measured essentially as previously described (*29*). Briefly, lung homogenates (1:10 in DMEM medium) were incubated with Vero E6 cells in 24-well culture plates for 40 min (37⁰ C 5% CO_2_), cells were washed 4 times with PBS and 1 time with DMEM medium and then incubated with DMEM medium (see above). Supernatant samples (200 µl) removed at 0, 24 and 48 hours were added to LBF lysis buffer (see above) and subjected to RT-PCR as described above.

### RNA-seq

RNA was isolated from lungs of mice using Qiagen RNeasy mini kits (Qiagen, CA, USA) with an on-column DNase step (Qiagen, CA, USA) according to the manufacturers’ instructions. RNA quantification was carried out in a Qubit fluorometer using the Qubit RNA HS assay kit (Invitogen, CA, USA).

RNA-seq was performed at the JP Sulzberger Columbia Genome Center (NY, NY). Libraries were generated using the Ilumina TrueSeq Standard mRNA kit according to manufacturers’ instructions. Polyadenylated RNA enrichment was performed. Sequencing of 100bp paired-end was performed on the Ilumina NovaSeq 6000 system. RNA-seq quality control was performed using fastQC v0.11.5, checking for per base sequence quality, per sequence quality scores and adapter content. Pseudoalignment to a kallisto index created from transcriptomes (Mouse: GRCm38) was performed using kallisto (0.44.0). We verified that each sample reached at least 75% of the target read goal and checked for adequate sequence alignment percentages. Sequencing yielded 19.5 M to 25.1M reads per sample resulting in identification of 35,199 transcripts. Analysis of differentially expressed genes under various conditions was performed using the R package DESeq2 v1/18.1, with default parameters. The mouse GRCm38 annotation file was downloaded from the ensemble BioMart website (https://www.ensembl.org/Mus_musculus/Info/Index). Gene Ontology (GO) enrichment analysis was carried out with Fisher’s exact test to estimate the specific GO categories, using the OmicsBox software (https://www.biobam.com/omicsbox).

### Histology

Lungs were rapidly isolated, carefully inflated and fixed in 4%-natural-buffered paraformaldehyde at room temperature (RT) for two weeks, followed by routine processing for paraffin embedding. Serial sections, 5 µm-thick, were cut and selected sections were stained with hematoxylin and eosin (H&E) for general histopathology and with Masson’s trichrome (MTC) for collagen and examined by light microscopy. Images were acquired using the Panoramic MIDI II slide scanner (3DHISTEC, Budapest, Hungary).

### Confocal microscopy

Lungs were processed as above and sections (5μm) were mounted on glass slides and deparaffinized. Antigen retrieval was performed by incubation in Target Retrieval Solution (DAKO, 30min, 95°C). After blocking in 5% BSA in PBS, slides were incubated (overnight, 4°C) with RBD-huFc fused protein (*18*) or with AChE-huFc fused protein (negative control, (*19*)). Anti-human Alexa Fluor 594-antibody was used for detection (Molecular probes®, ThermoFisher Scientific, Carlsbad, CA, USA). For nuclear staining, slides were mounted with Prolong® Gold antifade reagent containing DAPI (Molecular probes®, ThermoFisher Scientific, Carlsbad, CA, USA). Analysis was performed using an LSM 710 confocal scanning microscope (Zeiss, Jena, Germany) equipped with following lasers: argon multiline (458/488/514 nm), diode 405nm, DPSS 561 nm and helium-neon 633 nm. Fluorescence intensity was quantified using Zen software (version 2.1 Zeiss, Jena, Germany, 2008).

### Flow cytometry

Lungs harvested and cut into small pieces were digested (1h, 37c) with 4 mg/ml collagenase D (Roche, Mannheim, Germany). Tissue was meshed through a 40-µm cell strainer, and RBCs were lysed. Cells were stained using the following antibodies (eBioscience): CD45-FITC (clone 30-F11), CD31-PE (clone 390), CD326-PerCP (clone G8.8). Cells were defined as endothelial cells (CD45, CD31^+^) and epithelial cells (CD45, CD31^-^, CD362^+^). For ACE2 receptor staining, cells were stained using goat anti-mACE2 antibody (R&D, AF3437) followed by donkey anti-goat IgG coupled to AF647. For TMPRSS2 staining, cells were stained using anti-TMPRSS2 polyclonal antibody coupled to AF647 (Santa Cruz Biotechnology sc-515727). Cells were analyzed on FACSCalibur (BD Bioscience, San Jose, CA, USA).

### Statistical analysis

Simple comparisons were performed using the unpaired two-tailed Student’s *t*-test. Significance was set at *p*<0.05. Statistical analysis was calculated using Prism software (version 5.01, 2007; GraphPad Software, La Jolla, CA). All data are presented as means ± SEM.

## Results

### Induced pulmonary injuries predispose mice to SARS-Cov-2 infection

Cell entry of SARS-CoV-2 requires the binding of the viral spike to ACE2. Unlike ACE2 of humans, hamsters and ferrets, ACE2 of mice does not sensitize cells for infection (*12*). In line with this fact, CD-1 mice were refractive to SARS-CoV-2 infection as demonstrated by the lack of any discernable reduction in their body weights following viral infection (Fig. 1A). To determine whether pulmonary comorbidities affect the susceptibility of mice to SARS-COV-2, CD-1 mice were treated with one of three selected stimulants of acute lung injury, LPS, bleomycin or ricin. To promote the development of a mild and transient injury which resolves within a few days, these stimulants were administered intranasally at low doses, 1.7 mg/kg, 2U/kg and 1.7 μg/kg of LPS, bleomycin and ricin, respectively. The time-point at which the mice display an onset of disease as exemplified by body weight loss was 1, 4 and 2 days after the administration of LPS, bleomycin and ricin, respectively. At these time-points, the pre-administered mice were infected with SARS-CoV-2 at a dose of 5×10^6^ PFU/mouse and then monitored for 15 days. LPS-pretreated mice displayed an additional decrease in body weight following viral infection, however body weight regain began already at 5 dpi and the mice approached their weights determined prior to viral infection, within a short period of time (Fig. 1B). In contrast, mice pretreated with either bleomycin or ricin displayed a more sustained decline in body weight which lasted throughout the entire surveillance period following infection with SARS-CoV-2, reaching as low as 70% (bleomycin, Fig. 1C) and 77% (ricin, Fig. 1D) of their original weights determined immediately prior to viral infection. Most notably, 100 and 50 percent of the bleomycin- and ricin-pretreated mice, respectively, died within the 15 days surveillance period, death occurring at 4 to 7 dpi for bleomycin-pretreated mice and 9 to 13 dpi for ricin-pretreated mice (Fig. 1E).

**Fig. 1:**
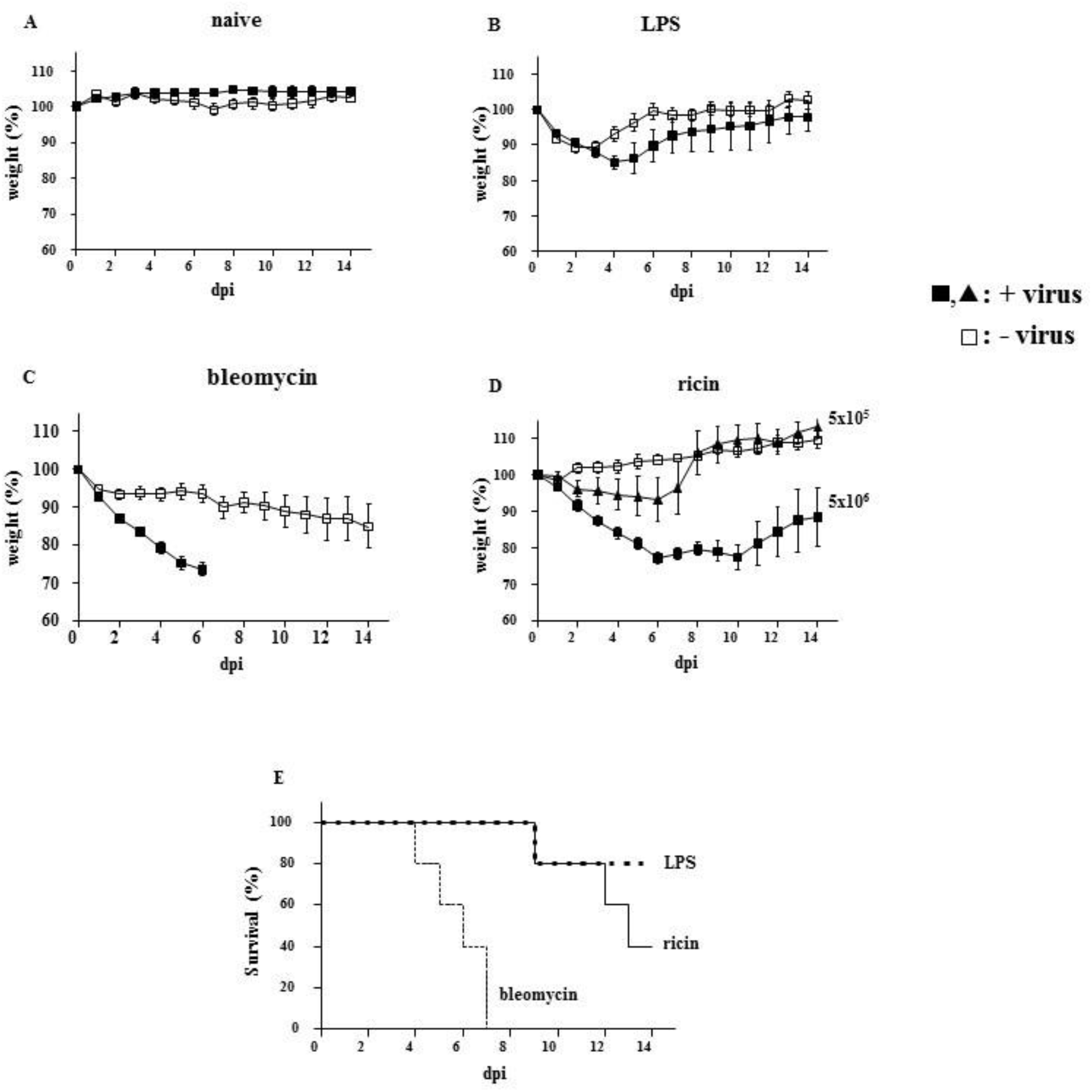
Effects of SARS-CoV-2 infection on body weight and mortality of CD-1 mice pretreated with ALI/ARDS stimulants. A-D: body weights of CD-1 mice (presented as percent of original weight determined at the time of viral instillation) were monitored over a period of 15 days following intranasal instillation of SARS-CoV-2 virus. A: Body weights of mice infected with virus at a dose of 5×10^6^ PFU per mouse (black squares) compared to body weights of naïve mice (white squares). B: Mice were administered LPS (1.7 mg/ kg body weight) and 1 day later were infected (black squares) or not (white squares) with virus (5×10^6^ PFU/mouse). C: Mice were administered bleomycin (2U/kg body weight) and 4 days later were infected (black squares) or not (white squares) with virus (5×10^6^ PFU/mouse). Due to significant mortality, only data of 0 to 6 days are presented for bleomycin-SARS-CoV-2 mice. D: Mice were administered ricin (1.7 μg/ kg body weight) and 2 days later were infected with virus at a dose of 5×10^5^ (black triangles) or 5×10^6^ (black squares) PFU/mouse or not (white squares). E: Kaplan-Meier survival curves of the mice groups exhibiting mortality: thick dashed line-LDR-SARS-CoV-2 (5×10^5^ PFU/mouse), black line-LDR-SARS-CoV-2 (5×10^6^ PFU/mouse), thin dashed line-bleomycin-SARS-CoV-2 (5×10^6^ PFU/mouse). Data represent mean ± SEM, n=5-6 per group.

The present study focused on the SARS-CoV-2 mice model predisposed by low dose ricin (LDR). Repeated experiments allowed us to determine that following infection with SARS-CoV-2 at a dose of 5×10^6^ PFU per mouse, death rates were within the range of 17-75% with an overall average of 39% (n=26, 5 independent experiments). When low-dose ricin (LDR) mice were subjected to SARS-CoV-2 infection at a lower viral dose, 5×10^5^ PFU per mouse, body weight reduction was less pronounced (<7%, Fig. 1D) and full weight regain was reached at 8 dpi. In a representative experiment (Fig. 1E), 20% of the animals died, similar to the average death rate determined in repeated experiments (n=16, 3 independent experiments, range=0-40%). Taken together, these findings clearly demonstrate that though CD-1 mice are refractive to SARS-CoV-2 infection, compromised pulmonary conditions induced by pre-exposure to low doses of selected ALI stimulants, confer a SARS-CoV-2-sensitive phenotype to these mice.

### Characterization of the pulmonary injury induced by intranasal application of low-dose-ricin

To characterize the impaired pulmonary background that promotes SARS-CoV-2 sensitivity, we monitored the pathological changes occurring in LDR mice. These examinations, which were carried out starting from the time of ricin application included surveillance of body weight and motor activity over time, differential blood counts and bronchoalveolar fluid (BALF) analyses. LDR mice which were not subjected to viral infection displayed a transient loss of weight of up to approximately 10% (9.6 ± 2.9%, Fig.2). In repeated experiments, maximal weight reduction occurred 2-6 days after administration of low dose ricin, while commencement of body weight regain was recorded between 3 to 7 days after ricin treatment. In most experiments, mice reached their initial body weights at days 7-12 and average body weight percentage at day 12 was 102.5±2.5%. LDR mice morbidity was also by following their motor activity, utilizing a recently developed computerized home cage monitoring system (HCMS100) based on laser-beam interruption counts (*13*). Examination of nocturnal activity profiles showed that LDR mice displayed a transient reduction in motor activity compared to sham-treated mice, which resolved at day 7 (Fig. 2).

**Fig. 2:**
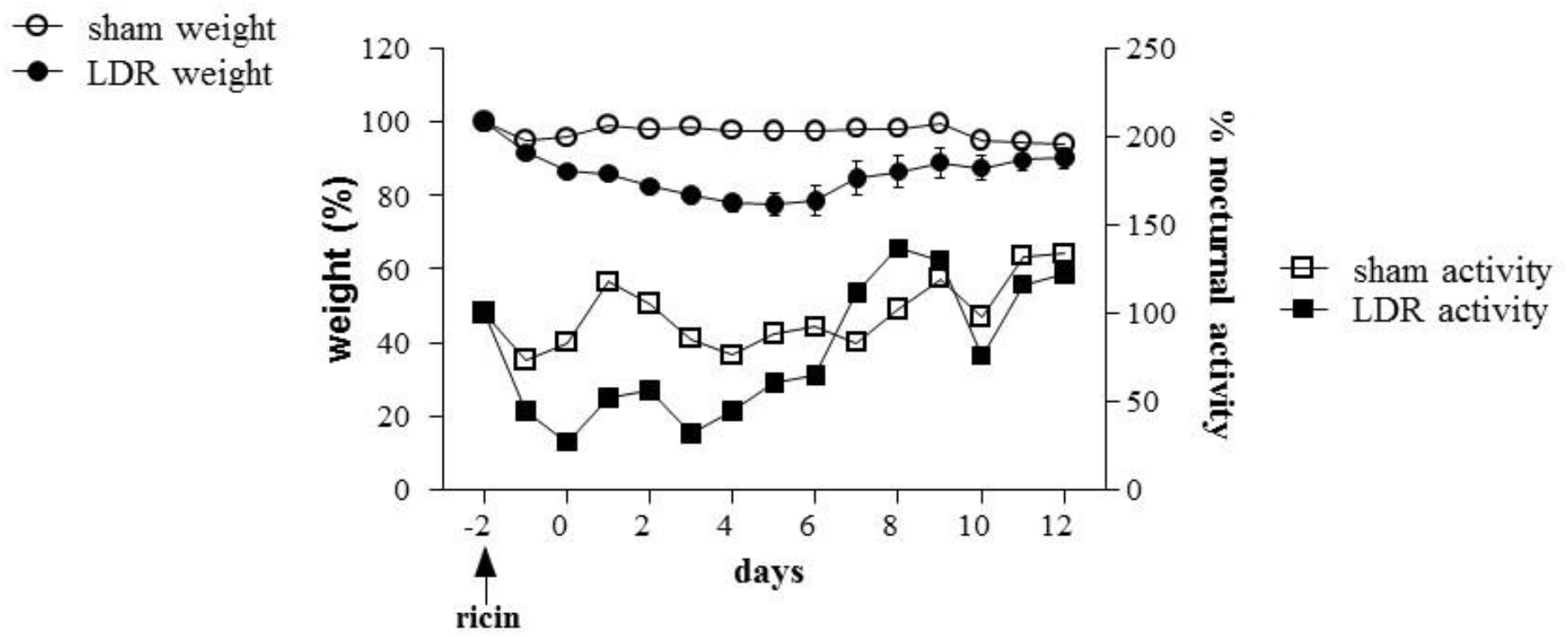
Effects of low dose ricin application on body weights and motor activity of CD-1 mice. CD-1 mice were monitored over a period of 15 days following intranasal application (indicated by arrow) of ricin (1.7 μg/kg) or PBS (sham treatment): circles, body weight of LDR (black)- and sham (white)-mice, squares: nocturnal motor activity profiles of LDR (black)- and sham (white)-mice. Body weights are expressed as percent of original weight determined at the time of ricin or sham administration and data represent mean ± SEM n=6; Nocturnal activity, group activity of 6 mice per home cage, statistics not provided.

Hematological analysis was performed on blood samples collected from LDR mice 2 days after the low dose ricin was administered (day 0), this time point corresponding to that at which LDR mice were infected with the virus. White blood cell counts in general and in neutrophils and monocytes in particular were elevated. Platelet counts increased as well (Table 1). BALF collected from LDR mice at this time-point displayed higher protein contents as well as elevated cell counts, compared to sham-treated mice. Elevated TNF-α, IL-1β and IL-6 levels were also measured in the BALF of the LDR mice (Table 1). In line with this observation, indicating that an inflammatory reaction has launched in response to low dose ricin administration, gene expression of cytokines and chemokines in the LDR mice lungs, 2 days after ricin application, were significantly upregulated, in comparison to naïve mice lungs (Fig. 3). Finally, assessment of ricin catalytic activity (28S rRNA depurination) in the lungs demonstrated that ∼8% of lung 28s rRNA extracted from LDR mice 2 days post-administration were depurinated at this time point (Table 1). At day 7, nearly all of the hematological and BALF-related parameters displayed values that did not differ significantly from the corresponding values determined in sham-treated mice (a single exception was the BALF protein concentration value which was significantly higher than in the sham mice). In addition, depurinated 28S rRNA, the hallmark of ricin catalytic activity, could not be measured above background level at day 7 (Table 1). Taken together, these observations suggest that the impaired pulmonary state in LDR mice is limited, resolution occurring within a matter of days.

**Table 1:** Blood counts, BALF analyses and ricin catalysis products in CD-1 mice following application of low-dose ricin.

|  |  | sham mice | LDR mice-day 0<br>(*p* value vs. sham) | LDR mice-day 7<br>(*p* value vs. sham) |
| --- | --- | --- | --- | --- |
| **complete blood count\*** | WBC (K/μl) | 4.51±1.68 | 10.64±1.78 (**0.003**) | 6.04±4.72 (0.564) |
| | NEU (K/μl) | 0.58± 0.92 | 1.82±4.81 (**0.007**) | 1.39±1.18 (0.496) |
| | MO (K/μl) | 0.17±0.17 | 0.73±0.22 (**0.007**) | 0.14±0.14 (0.809) |
| | PLT (K/μl) | 499±220 | 942±197 (**0.024**) | 734±217 (0.205) |
| **bronchoalveolar lavage fluid \*\*** | protein (mg/ml) | 1.56±0.21 | 2.01±0.18 (**0.006**) | 4.44±1.7 (**0.007**) |
| | cells ($\times 10^6$) | 36±27 | 86±27 (**0.019**) | 17.2±7.6 (0.178) |
| | IL-6 (pg/ml) | 34±29 | 586±110 (**4.47E-06**) | 75±30 (0.06) |
| | IL-1β (pg/ml) | 1±2 | 28±15 (**0.004**) | 43±41 (0.052) |
| | TNF-α (ng/ml) | 8±12 | 38±12 (**0.005**) | 28±22 (0.109) |
| **ricin catalysis products in lungs\*\*** | depurinated 28S rRNA (%) | 0.22±0.31 | 7.94±3.11 (**0.00056**) | 0.42±0.6 (0.53) |

\* n=4; \*\* n=5. Statistically significant *t*-test values (*p*<0.05) are presented in bold type.

**Fig. 3:**
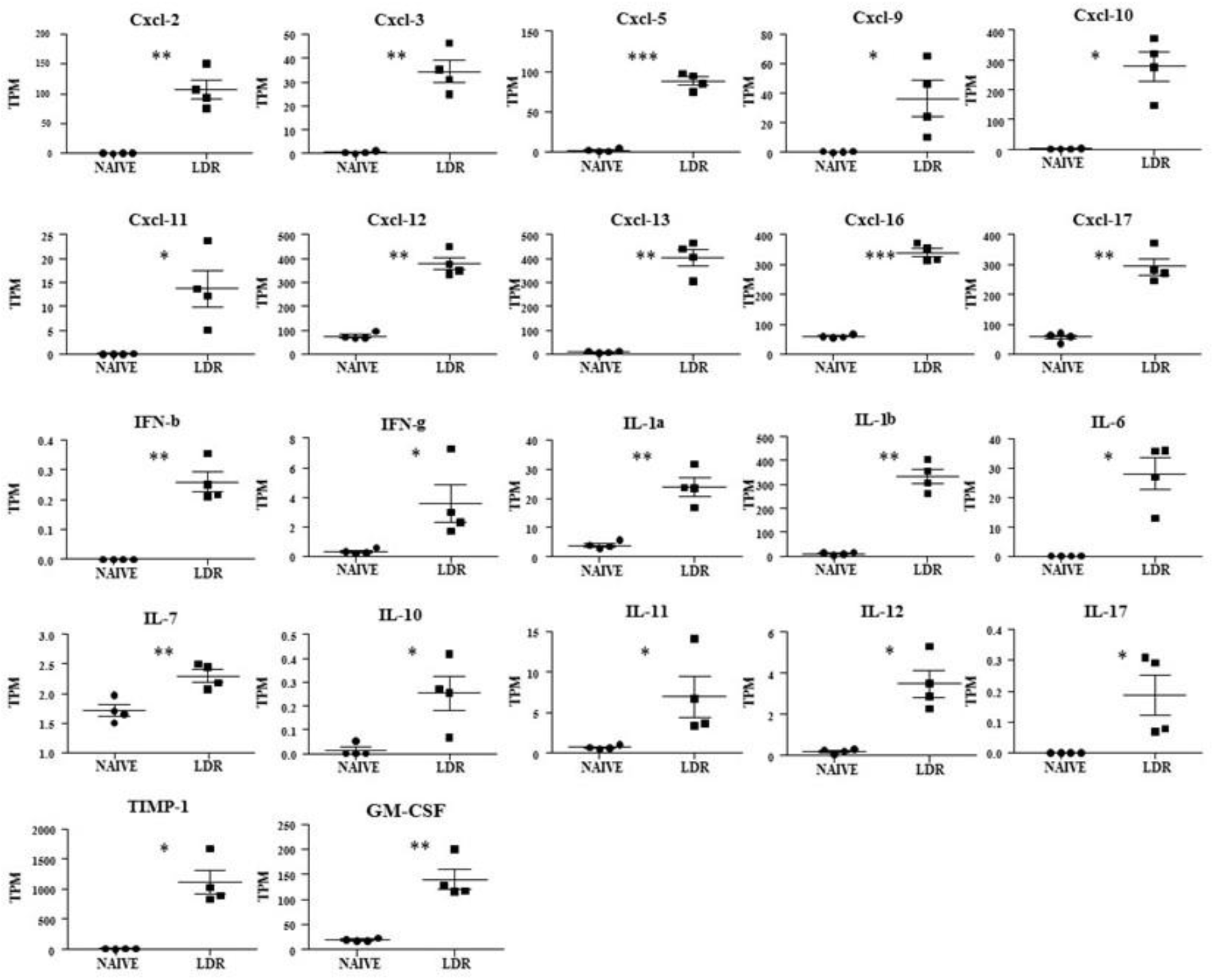
Cytokine and chemokine gene expression in lungs of LDR mice. Normalized gene expressed levels (transcripts per million-TPM) of 22 indicated cytokines and chemokines determined from RNAseq data of LDR mice (2 days post-low-dose-ricin-application) compared with naïve control. Data are represented as average ± SEM (n=4 per group). \**p* <0.05, \*\**p* <0.01, \*\*\**p* <0.001

To provide a more comprehensive view of the alterations in protein expression that might play a role in sensitizing LDR mice to SARS-CoV-2 infection, whole lung cell RNA-seq of LDR-*versus* naïve-mice were performed (Fig. 4). Principal component analysis revealed distinct transcriptional signatures between naïve and LDR mice (Fig. 4a). Compared with the naïve group, there were 8545 differentially expressed genes (DEGs) in the LDR group (2 days after low-dose-ricin application), comprising 4394 upregulated and 4151 downregulated genes (Fig. 4b). Gene Ontology (GO) analysis ranked by the false discovery rate (q value) after eliminating redundant terms, revealed 1379 G0 terms in upregulated DEGs (biological process-865, cellular function-210, molecular function-304) and 1289 GO terms in downregulated DEGs (biological process-810, cellular function-160, molecular function-319). These results reflect the notable magnitude of alterations that occur in the transcriptome profile of the LDR mice in a wide range of genes and in all three domains of the gene annotation analysis. Observing the top 50 (displaying highest q values, see Fig. S1) up-regulated biological processes GO terms, have shown an over-representation of themes related to immune and inflammatory response (44%) (Fig. 4c), corroborating our observation that low dose ricin application stimulates expression of proinflammatory gene pathways even though it does not cause a clinically acute disease.

**Fig. 4:**
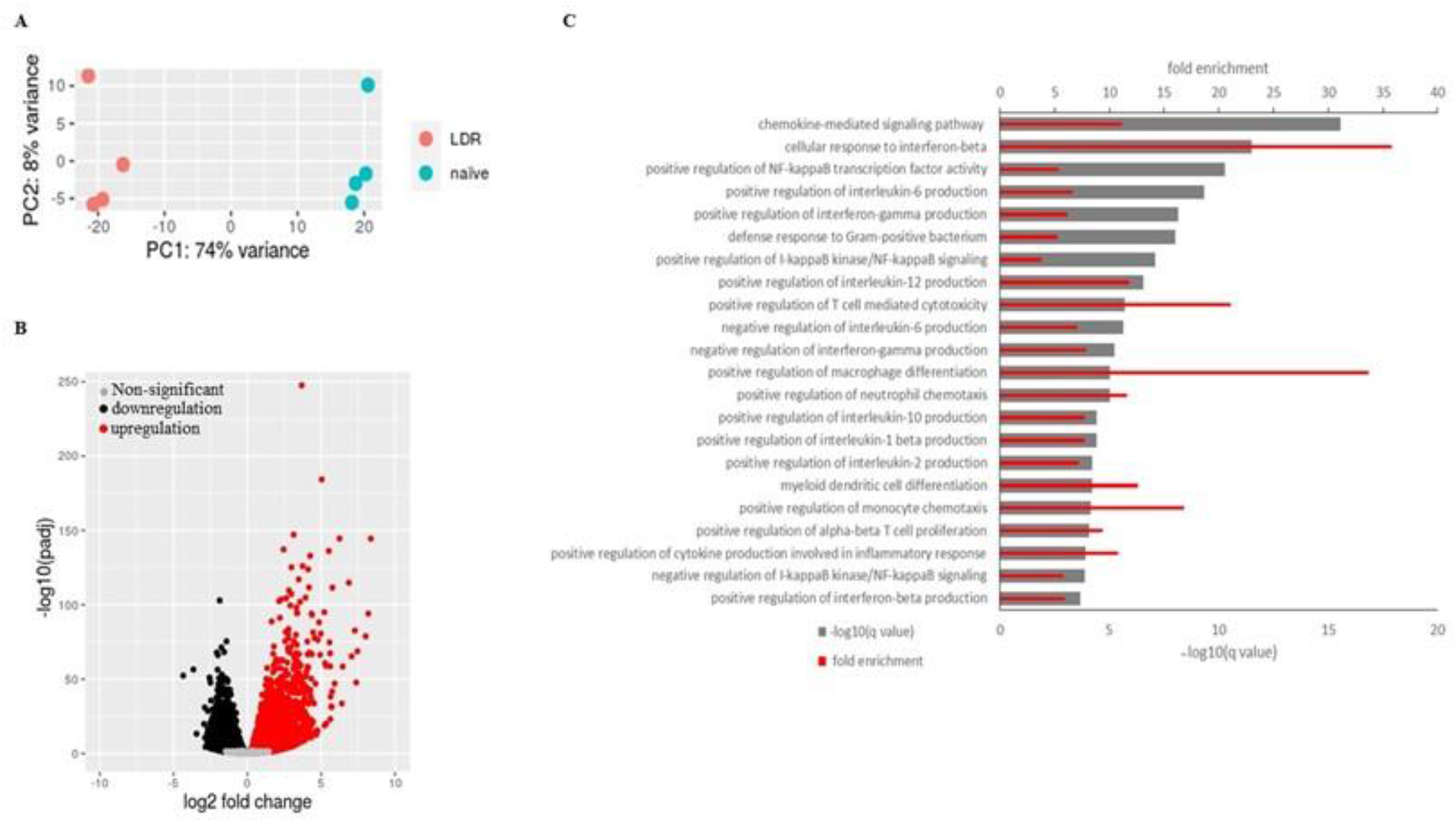
Distinct transcriptional signatures in LDR mice compared to naïve mice. RNAseq analysis of lung homogenates from naïve- and LDR-(2 days post-low-dose-ricin application) mice (n=4). A: Principal component analysis was performed for 8 samples with log2 transformed gene-level normalized counts data. PC, principal component. B: Volcano plot comparing differentially expressed genes, respectively, with a false discovery rate (q value) < 0.05. Multiple comparisons were accounted for by calculation of a Benjamini-Hochberg false discovery rate-adjusted P value (q value). Each dot in the volcano plot represents a single gene. C: Gene Ontology (GO) enrichment analysis of biological process (BP) terms enriched in upregulated genes from comparisons of LDR mice versus naïve. Terms were ranked by the false discovery rate (q value) and the immune and inflammatory terms are listed from the top 50 after eliminating redundant terms. Gray bars show the –log10[q value]. Red bars show the genes fold enrichment.

We further ascertained that the marked deterioration observed in LDR-SARS-CoV-2 mice is directly due to the viral infection and does not reflect an exacerbation of a ricin-related disease. To this end, LDR-SARS-CoV-2 mice were treated 2 hours prior to SARS-CoV-2 infection with equine-derived anti-ricin F(ab’)_2_ antibodies at a >10-fold higher dose than that required for full protection of mice from a lethal dose (7 μg/kg body weight) of ricin (*14, 15*). Indeed, treatment of mice with these antibodies at the time of low dose ricin treatment prevented body weight loss (data not shown), attesting to efficient and rapid neutralization of the toxin. Nevertheless, treatment with anti-ricin antibodies did not have any measurable beneficent effect on the well-being of the LDR-SARS-CoV-2 mice; following viral infection, weight loss continued to decline markedly (Fig. 5) in a manner similar to that observed in LDR-SARS-CoV-2 mice that were not treated with anti-ricin antibodies (Fig. 1D). Twenty percent of the anti-ricin-antibody-treated LDR-SARS-CoV-2 died within the 15 day surveillance period, this death rate falling within the death rate range documented for mice that were not subjected to anti-ricin antibody treatment. Thus, the significant signs of morbidity documented in LDR mice infected with SARS-CoV-2 virus stem from the viral infection and not from continuous ricin toxic activity.

**Fig. 5:**
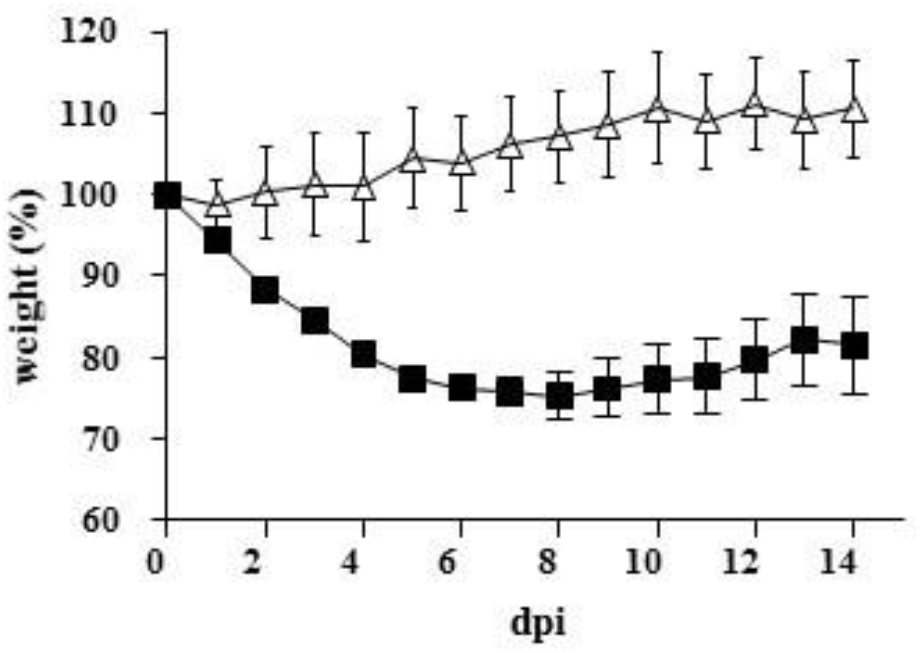
Anti-ricin treatment does not alleviate SARS-CoV-2-induced body weight loss of LDR mice. Two days after intranasal application of low-dose ricin (1.7 μg/kg), mice were treated (i.p.) with anti-ricin antibodies (1730 IsNU/mouse, (*12*)) and 2 hours later were infected (black squares) or not (white triangles) with SARS-CoV-2 virus (5×10^6^ PFU/mouse). Mice body weights of CD-1 mice were monitored over a period of 15 days following viral infection. Data, presented as percent of original weight determined at the time of virus instillation, represent mean ± SEM, n= 4-5 per group.

### Characterization of the SARS-COV-2 infection model in LDR mice

We evaluated the impact of SARS-CoV-2 infection on lung histopathology using H&E and MTC stainings (Fig. 6). At 7 dpi, lungs from LDR-SARS-CoV-2 mice exhibited severe damage manifested by extensive peribronchial and perivascular inflammatory cell infiltration along with intra-alveolar edema, fibrin and macrophage accumulation (Fig. 6, B, C, right). In contrast, lungs from SARS-CoV-2 mice (Fig. 6, A, right) exhibited a relatively preserved alveolar structure and a mild immune peribronchial and perivascular cell infiltration. Upon viral infection (Fig. 6, day 0), LDR mice exhibited mild mononuclear cell infiltrations in the peribronchial and perivascular areas (Fig. 6, B, C, left). Taken together, these observations show that severe manifestations of SARS-CoV-2 induced lung injury were restricted to LDR mice.

**Fig. 6:**
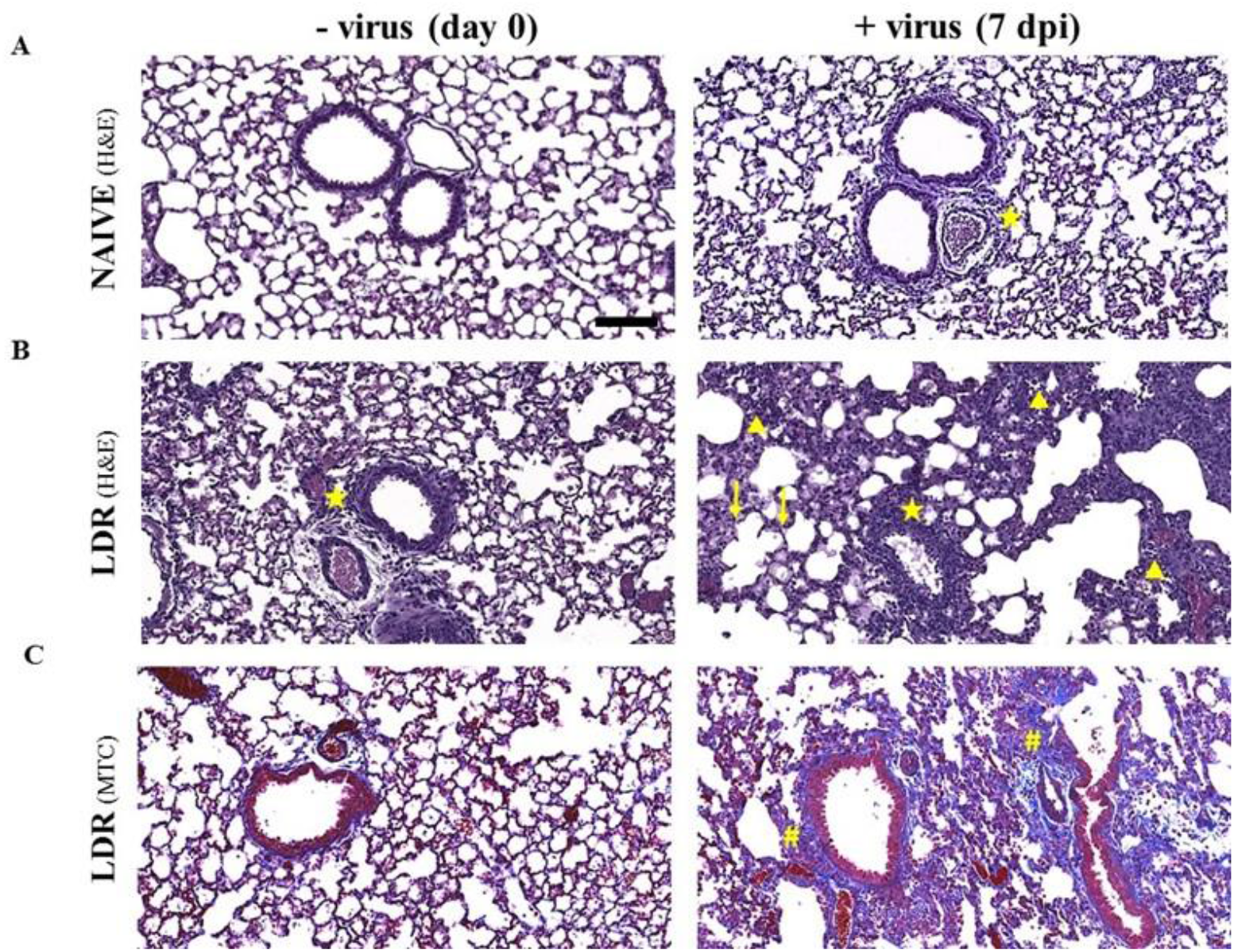
Histology of naïve and LDR mice lungs before and after SARS-CoV-2 infection. A: naïve and B-C: LDR mice (ricin application at day -2) were subjected to SARS-CoV-2 infection at a dose of 5×10^6^ PFU/mouse. Lungs at the time of viral infection (day 0) and 7 days later (7 dpi) were harvested, fixed and processed for paraffin embedding. Sections (5 µm) were stained with H&E (A, B) or MTC (C). Representative sections of n=4 are shown. Indicated are: perivascular and peribronchial inflammation (stars), infiltration in perivascular and alveolar sites (triangles), edema (arrows) and fibrin (hashtags). Magnification X20, bar=100 µm.

To further characterize the effect of low dose ricin administration on SARS-CoV-2 infectability, we exposed naive and LDR mice to SARS-CoV-2 at a dose of 5×10^6^ PFU per mouse and thereafter, various organs and tissues were harvested and examined for the presence of viral RNA. Already at 2 days after viral infection, viral RNA levels in nasal turbinate, trachea and lung homogenates were markedly higher in LDR-SARS-CoV-2 mice than in SARS-CoV-2 mice (Fig. 7A). Significantly higher levels of viral RNA were also documented as early as 2 days post-infection in the serum and heart of LDR-SARS-CoV-2 mice (Fig. 7B). Higher levels of viral RNA were also detected in brain and spleen homogenates of LDR-SARS-CoV-2 mice, however these did not differ statistically from the corresponding levels determined in SARS-CoV-2 mice. No viral RNA was found in liver and kidney homogenates (Fig. 6B). We further profiled viral RNA in serum and heart homogenates over time. Indeed, LDR-SARS-CoV-2 mice displayed significantly high levels of viral RNA in the serum and heart at all time-points examined, 3, 5 and 7 dpi (Fig. 7C). Examination of lung homogenates at these time-points, demonstrated a progressive decline of viral RNA over time, yet levels remained relatively high even in SARS-CoV-2 mice (Fig. 7C). To determine the presence of viable SARS-CoV-2, lung homogenate samples were subjected to growth kinetics profiling. To this end, Vero E6 cells were with incubated with homogenate lung samples harvested at 3 dpi, and extracellular viral RNA was repeatedly quantified by RTPCR at 3 different time-points. Unlike lung homogenates of SARS-CoV-2 mice which exhibited unaltered Ct values, those of LDR-SARS-CoV-2 mice displayed decreasing Ct values of 29, 23 and 8 at 0, 24 and 48 hours, respectively (Fig. 7D), attesting to the fact that viable virus is present at 3 dpi, solely in LDR-SARS-CoV-2 mice.

**Fig. 7:**
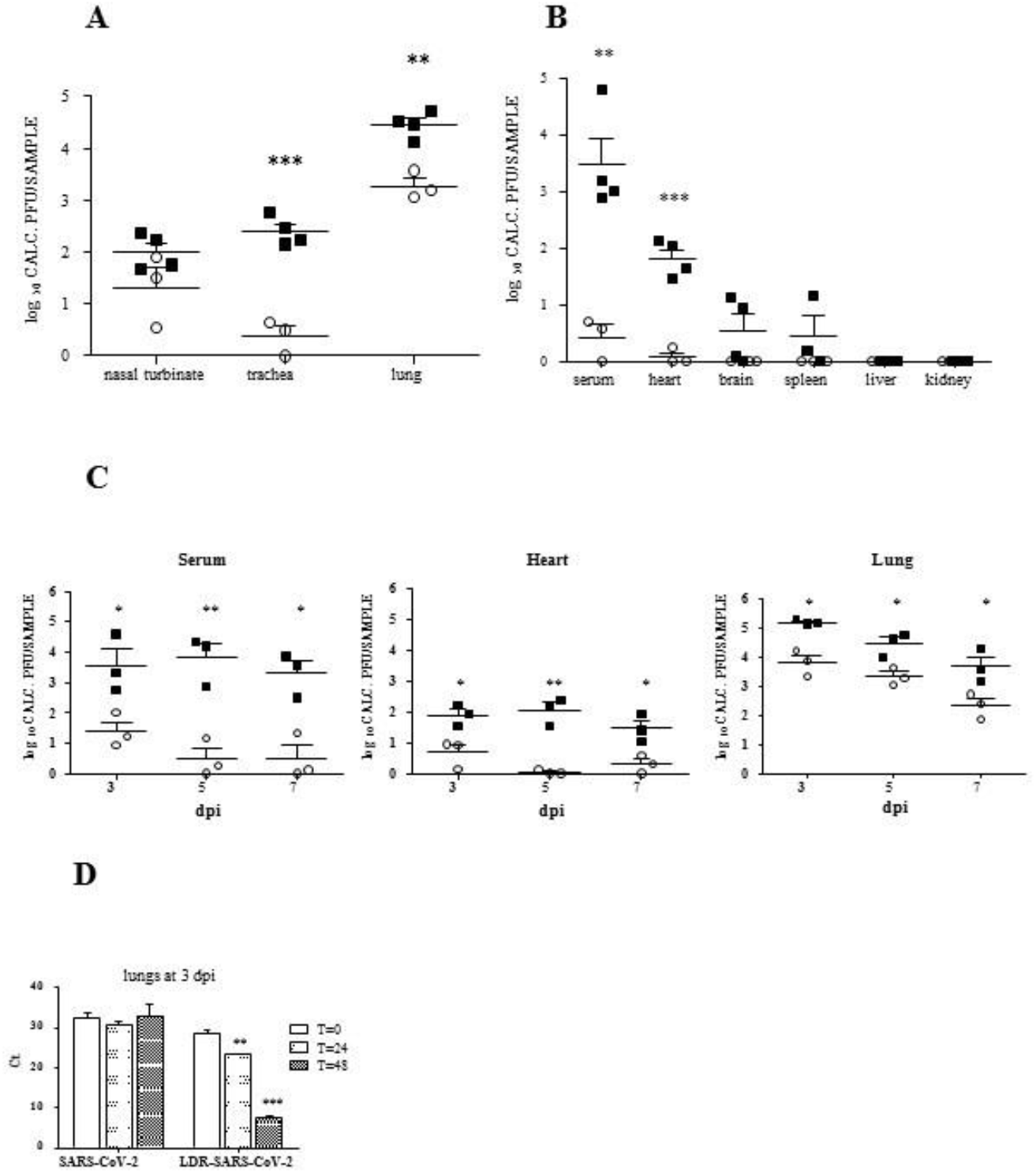
Viral RNA levels in various tissues derived from SARS-CoV-2 and LDR-SARS-CoV-2 mice. A-C: Viral loads were determined by RT-PCR in homogenates prepared from the indicated tissues of SARS-CoV-2 mice (empty circles) and LDR-SARS-CoV-2 mice (filled squares). A PFU calibration curve tested in parallel was utilized to express Ct values as calculated PFUs. A-B: viral loads determined at 2 dpi. C: viral loads determined at 3, 5 and 7 dpi. D: SARS-CoV-2 growth kinetics in lung homogenates prepared at 3 dpi was performed by RT-PCR at the indicated time-points. Data represent mean ± SEM, n= 3-4 (A-C) or 2-3 (D) per group. \**p* <0.05, \*\**p* <0.01, *** *p* <0.001.

### Anti-viral-related antibodies protect LDR mice from SARS-COV-2 infection

If indeed the deleterious manifestations documented in LDR-SARS-CoV-2 mice are due to direct viral activity, one may expect them to be redressed by treating the mice with anti-SARS-CoV-2-related antibodies. To address this issue, LDR mice were treated with polyclonal antibodies raised against either the entire virus or the receptor binding domain (RBD) portion of the SARS-CoV-2 spike (see Materials & Methods) or with the human MD65 monoclonal antibody shown to target the SARS-CoV-2 RBD (*13*). Application of either of the three anti-SARS-CoV-2-related antibodies 2 hours prior to viral infection, promoted mouse body weight regain (Fig. 8). Accordingly, at 9-10 dpi the body weights of the mice groups treated with each of the three antibody preparations approached their weight values determined at the time of SARS-CoV-2 infection. No deaths were recorded in the mice groups treated with either of the anti-viral related antibodies. In contrast, mice treated with non-specific antibodies (normal rabbit serum or TL1 Mab isotype control, (*16*)) displayed a steady drop in body weight and 40-100% of the mice succumbed within 5-13 dpi.

**Fig. 8:**
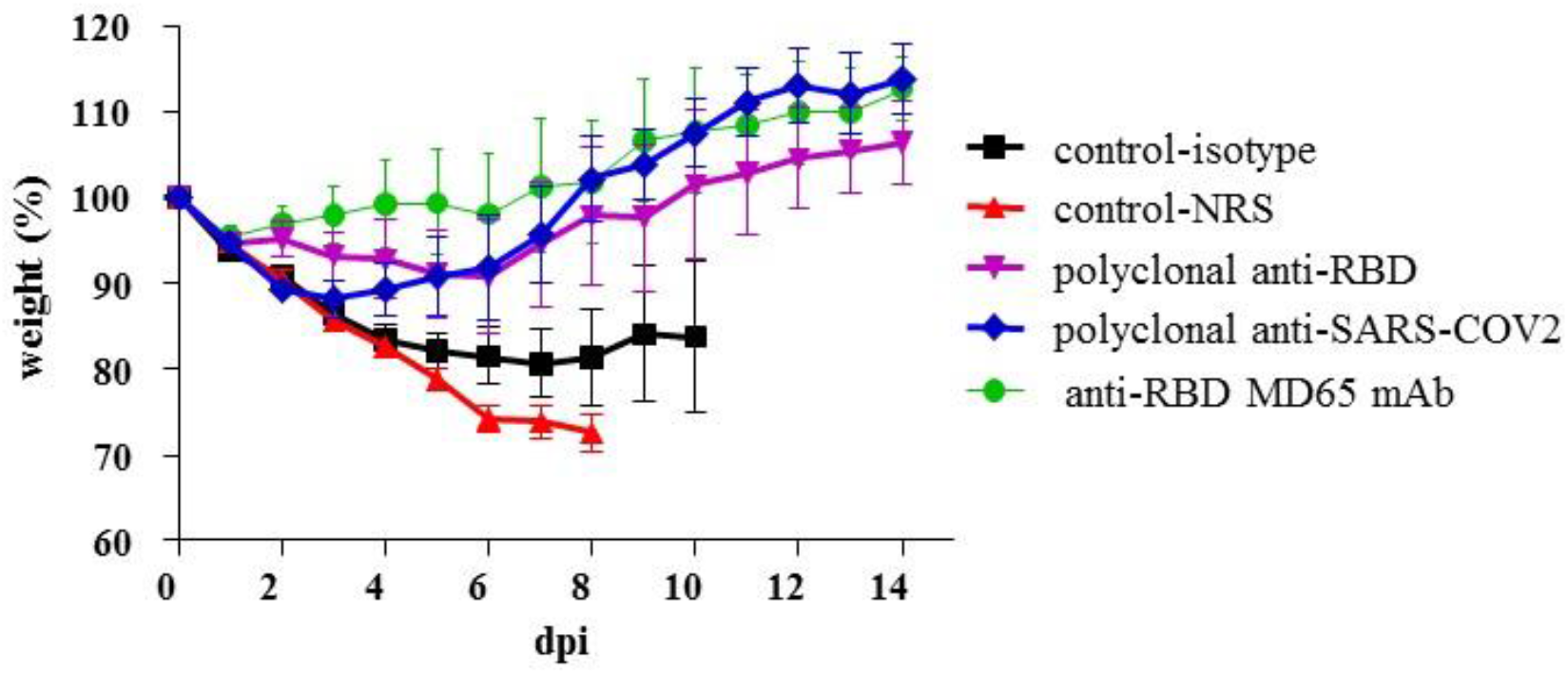
Anti-SARS-CoV-2-related antibodies protect LDR mice from SARS-CoV-2 infection. Two days after intranasal application of low-dose ricin (1.7 μg/kg), mice were treated (1 ml/mouse, i.p.) with the following SARS-CoV-2-related antibodies: rabbit-derived polyclonal anti-SARS-CoV-2 antibodies (blue diamonds), rabbit-derived polyclonal anti-RBD antibodies (magenta triangles) and anti-RBD MD65 mAb (green circles). Non-related isotype control (black squares) and normal rabbit serum (NRS, red triangles) served as negative controls for the SARS-CoV2-specific monoclonal and polyclonal antibodies, respectively. Mice body weights were monitored over a period of 15 days following viral infection. Due to significant mortality, only data of 0 to 8-10 days are presented for the control groups. Data, presented as percent of original weight determined at the time of virus instillation, represent mean ± SEM, n= 4-5 per group.

### Effect of LDR pretreatment on ACE2 and TMPRSS2 expression

ACE2 is the key cell-surface receptor for SARS-CoV-2 while the TMPRSS2 protease serves as a co-factor involved in the trimming of the viral spike, a process which facilitates viral internalization (*17*). It has been well established by others and shown in the present study as well, that mice are not susceptible to SARS-CoV-2 infection and as such, cannot serve as models for the CoVID-19 disease. This refractivity towards the virus is attributed to the inability of SARS-CoV-2 to bind effectively to murine ACE2 (*12*). Yet, in view of the fact that low doses of selected ALI stimulants render mice sensitive to the virus, we examined whether application of these stimulants affected the expression of ACE2 and TMPRSS2. To this end, lungs of LDR mice were harvested 2 days after application of ricin. This time point was chosen since alterations documented at this time-point, correspond to those which occur at the time of viral infection following treatment with low-dose-ricin. Cell suspensions prepared from the mice lungs were analyzed by flow cytometry, utilizing anti-mACE2 and anti-TMPRSS2 antibodies (Fig. 9 and Fig. S2).

**Fig. 9:**
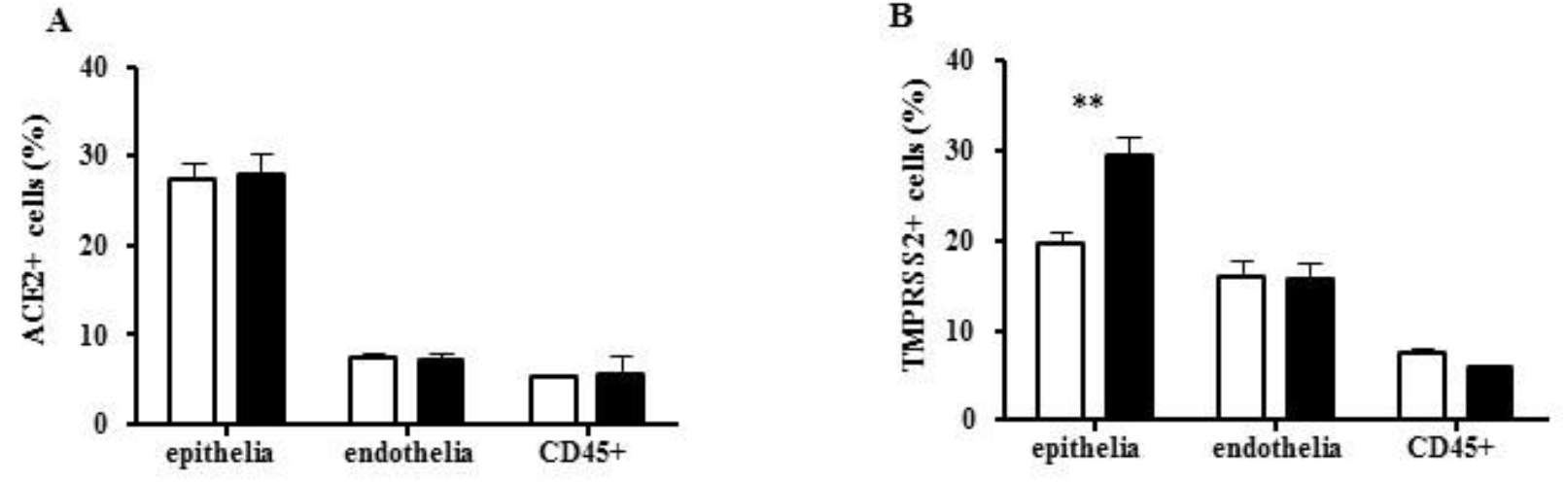
Expression of ACE2 and TMPRSS2 in lung cell types of LDR mice. Lungs harvested from LDR (filled bars) and sham-treated (empty bars) mice 2 days after treatment, were analyzed by flow cytometry for (A) ACE2 (n=3) and (B) TMPRSS2 (n=6). Data, referring to percent of positive cells out of the indicated cell-types, are presented as mean ± SEM, \*\**p* <0.005.

Comparison between sham-treated and LDR mice demonstrated that ricin administration did not bring about any noticeable increase in the number of cells expressing ACE2 (Fig. 9A). In contrast, the number of lung cells expressing TMPRSS2 was significantly higher in LDR mice than in the sham-treated mice (Fig. 9B). Notably, this increase in TMPRSS2 expression was cell-type dependent. Thus, a 50 percent increase was observed in the number of TMPRSS2^+^ lung cells of epithelial (CD45^-^CD31) lineage, while lung endothelial (CD45^-^CD31^+^) and hematopoietic cell (CD45^+^) TMPRSS2^+^ counts (Fig. 9B) were not affected.

The fact that the number of cells expressing cell-bound ACE2 did not increase following treatment with ricin, does not negate the possibility that viral entry into lung cells of LDR mice takes place by the binding of SARS-CoV-2 spike RBD to the murine ACE2 with the aid of factor(s) which are present only following induction of the pulmonary injury or by binding to cell-surface receptors other than ACE2 which are newly expressed/exposed following mice sensitization. To address this issue, we examined whether application of low-dose-ricin affects binding of RBD to mice lung cells. To this end, lung sections prepared from LDR- and naïve-mice, were incubated with RBD-huFc fused protein (*18*) or with AChE-huFc (used as control for Fc) fused protein (*19*), immunostained with AF594-labeled anti-huFc antibodies and visualized by confocal microscopy (Fig. 10). Lungs of LDR mice displayed markedly higher binding of RBD-huFc than those of naïve mice (Fig. 10A). This staining reflects authentic RBD binding to the lung cells and not huFc binding, as incubation with control AChE-huFc-fused protein resulted in no noticeable binding (LDR-Fc control). Quantitation of the mean fluorescence intensity in each scanned field (5 fields/lung) demonstrates a 70% increase RBD binding in LDR mice lungs (Fig, 10B). Whether this increased binding of RBD to LDR lungs is clinically functional and relevant, needs to be further examined.

**Fig. 10:**
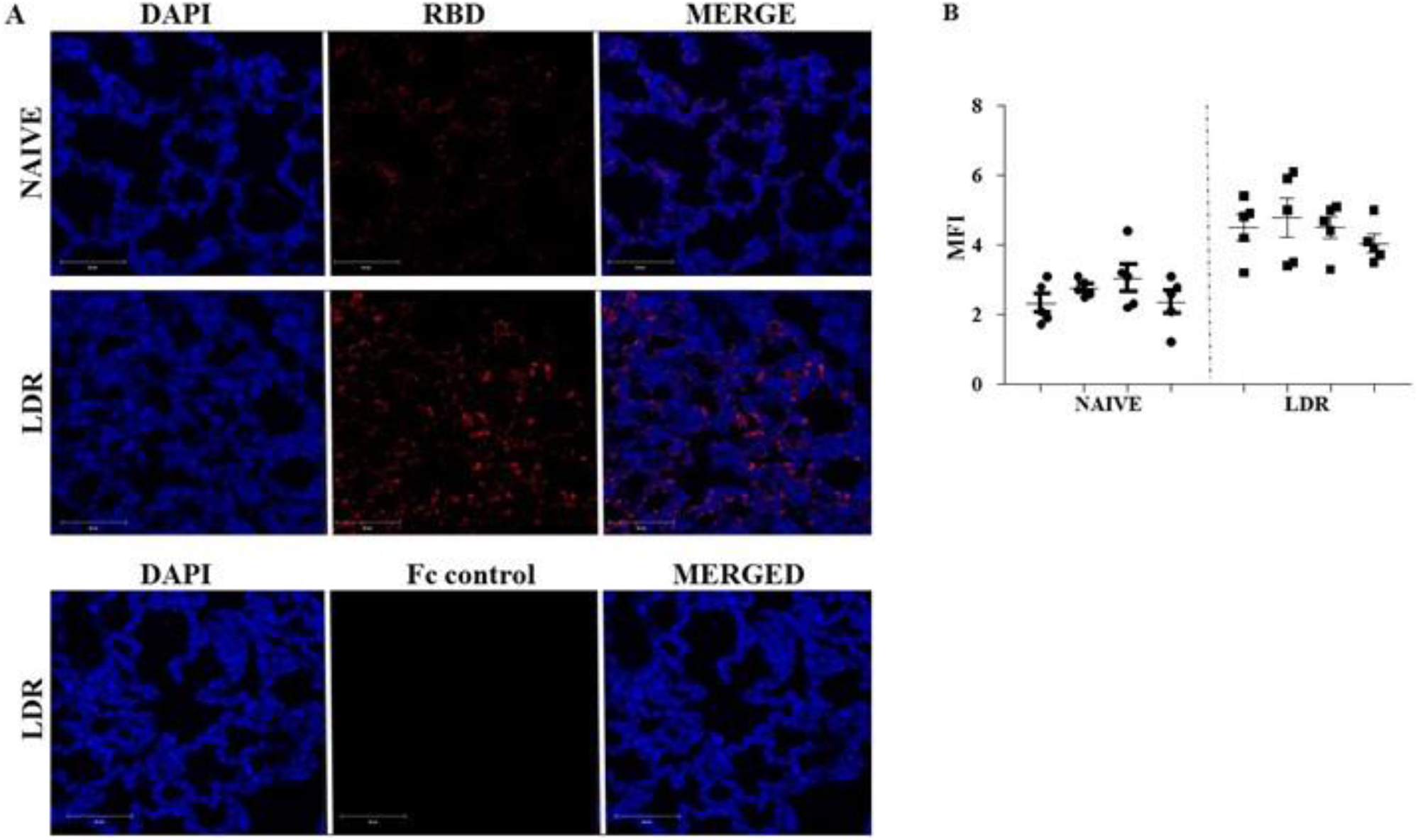
RBD binding to lung sections of LDR mice. **A**. Confocal microscopy scans of lung sections. Sections from LDR (2 days after low-dose-ricin administration) and naïve mice were incubated with RBD-huFc or AChE-huFc (Fc control) and then immunostained with anti-human AF594. Staining of RBD in red and identification of nuclei by DAPI in blue. **B**. Scatterplots of the fluorescence staining intensities of RBD expressed as mean fluorescence intensity (MFI) (n=4 mice, 5 scanned fields/lung).

## Discussion

In the present study we show that transient pulmonary insults prompted by administration of low doses of the ALI/ARDS stimulants, LPS, bleomycin or ricin, render mice to be sensitive to SARS-CoV-2 viral infection. Recently, mice models were adapted for SARS-CoV-2 infection either by adjusting the viral spike RBD domain to fit the murine version of ACE2 (*20, 21*) or by transient or constitutive expression of the human ACE2 in mice (*7, 8*). In contrast, the model presented in the current study consists of mice expressing their native ACE2 receptor yet are sensitive to genetically unaltered *bona fide* SARS-CoV-2 virus.

When LPS-treated mice were subjected to SARS-CoV-2 infection, the deleterious effect of the virus, reflected by additional loss of body weight, was slight and the mice displayed body weight regain within a few days. This narrow dynamic range of measurable viral-induced body weight loss precludes the use of LPS-pretreated mice for meaningful evaluation of virus-induced pathologies or anti-viral medical countermeasures. On the other hand, following SARS-CoV-2 infection of ricin- or bleomycin-pretreated mice, body weights of the mice decreased steadily and significantly during the entire course of the experimental surveillance period (15 days). Most importantly, 50-100 percent of the ricin- and bleomycin-pretreated mice animals died following viral exposure.

The finding that SARS-CoV-2 sensitivity is retained even when the LDR mice were treated with anti-ricin antibodies before viral infection, implies that active ricin is not required for conferring sensitivity towards the virus. Rather, the low-dose-ricin-induced pulmonary pathologies themselves promote viral susceptibility in these mice. Pathophysiological assessment of LDR mice at the time of viral infection, identified various factors that may contribute to the pathological setting required for acquisition of sensitivity towards SARS-CoV-2. These include high-level expression of proinflammatory cytokines and chemokines in the lungs and serum, mild immune cell infiltration to the lungs and elevated protein levels in the lungs, the latter indicative of compromised alveolar-capillary barrier function. Factors that may bring about gain-of-sensitivity towards SARS-CoV-2 in bleomycin-treated mice were not addressed in the present study and we now plan to examine these in our laboratory and to determine whether some of these overlap with the factors defined for low-dose-ricin-pretreated mice.

Quantitative RT-PCR analysis of tissues derived from LDR-SARS-CoV-2 and SARS-CoV-2 mice, demonstrated the presence of significantly higher levels of viral RNA in the lungs of LDR-SARS-CoV-2 mice. High levels of viral RNA, which were detected in the hearts and serum of LDR-SARS-CoV-2 mice, remained high at all examined time-points. Conversely, viral RNA levels in the hearts and serum of SARS-CoV-2 mice were considerably lower and waned rapidly. A recently published study also reported the presence of extrapulmonary SARS-CoV-2 virus in organs such as the heart, in a BALB-C-based mice model infected by murine-adapted SARS-CoV-2 virus (*21*). This finding is of considerable clinical relevance since autopsy samples obtained from COVID-19 patients revealed the presence of the virus in organs other than the lungs, including the heart, brain and blood (*22*). In some cases, brain and heart damage were also documented in COVID-19 patients (23) yet whether these injuries stem from direct viral activity or are due to an excessive immune response and virus-induced cytokine storm remains to be elucidated. Studies at our laboratory are now being performed to determine the presence and extent of organ injury in the hearts and brains of LDR-SARS-CoV-2 mice.

Taken together, our findings raise the intriguing question as to how the virus gains entrance into cells of CD-1 mice. Protection of LDR-SARS-CoV-2 mice by application of anti-RBD antibodies seems to imply that viral entry into cells of low dose ricin-treated mice is mediated *via* the RBD region of the viral spike. One may therefore be tempted to suggest that increased expression of ACE2 can compensate for the relatively poor binding of SARS-CoV-2 spike RBD to this ACE2 species and that overexpression of this receptor in mouse lung cells can overcome this limitation and bring upon a successful viral infection. However, comparative flow cytometry analysis of lung cells derived from sham-treated and LDR mice failed to detect an increase in ACE2 expressing cells following application of low dose ricin. On the other hand, histochemical analysis of lung sections prepared from LDR mice did reveal a significant increase in the binding of viral RBD. The combination of these two findings may imply that in LDR mice (and possibly in bleomycin mice), there is a gain of receptors other than ACE2 that can serve as an effective binding site for SARS-CoV-2, perhaps with the aid of the high level of lung cell surface TMPRSS2 induced by low dose ricin application.

Various viruses use junction proteins as receptors and access epithelium cells through apical junction complexes which contain tight junctions (TJs) and adherens junctions (AJs) even if most of these proteins are not readily available, possibly by taking advantage of compromised epithelium (*24*). Indeed, studies carried out in our laboratory demonstrated that intranasal application of a lethal dose of ricin (7μg/kg) to mice leads to rapid diminution of both adherens and tight junctions in the lungs, thereby precipitating the disruption of the alveolar-capillary barrier (*25*). Intriguingly, altered expression of tight junction molecules in the alveolar septa has also been reported in bleomycin- and LPS-induced lung injury models (*26, 27*). In line with the notion that alterations in junction proteins integrity play a role in the acquired sensitivity to SARS-CoV-2, Gene Ontology analysis of the RNA-seq upregulated genes in lungs of LDR mice at the time of viral infection, demonstrated, amongst others, the enrichment of pathways involved in protein binding, including enrichment of gene clusters involved in cadherin binding pathway (molecular function in Supplement 1).

Whether aberrations of junction proteins indeed play a significant role in the sensitization of mice towards SARS-CoV-2 remains to be determined. Nevertheless, and irrespective of the exact mechanism by which SARS-CoV-2 virus gains entrance to CD-1 mice lung cells, the existence of a second tier of entry points for SARS-CoV-2 in the case of pulmonary comorbidities, may have important implications on COVID-19 in humans, especially in high-risk populations. Decoding the process by which SARS-CoV-2 gains entrance into murine cells following sensitization by ricin or bleomycin is therefore a focal point of the ongoing research at our laboratory.

In summary, we established a unique mouse model for the study of the molecular pathways involved in comorbidity-dependent SARS-CoV-2-induced pathologies and for examining potential medical countermeasures for treatment of COVID-19 in high risk populations.

## General

We thank Drs. Ziv Oren, Adi Beth-Din and Anat Zvi for fruitful discussions and suggestions.

## Author contributions

Conceived and designed the experiments: R.F., C.K., T.S. and S.Y.

Investigation: R.F., L.B.-O., S.L., T.K., H.G, M.A., D.G., Y.E., E.M., I.C.-G., G.Z., O.S., O.I., D.S., O.M., E.Z., Y.V., S.M., B.P., H.A., C.K. and T.S.

Resources: H.L., O.M., R.R., R.A., T.I., S.M., H.A., I.G., A.B.-S.

Writing original draft: C.K. and T.S.

Reviewing: R.F., O.M., L.B.-O., S.L., T.K., T.I., S.M., B.P., H.A., I.C.-G., R.R.

## Data and materials availability

The transcriptomic data have been deposited to the NCBI database (GEO accession number of the transcriptome series is GSE159461 and the SRA BioProject number is PRJNA669006).

## Supplementary Materials

**Fig. S1:**
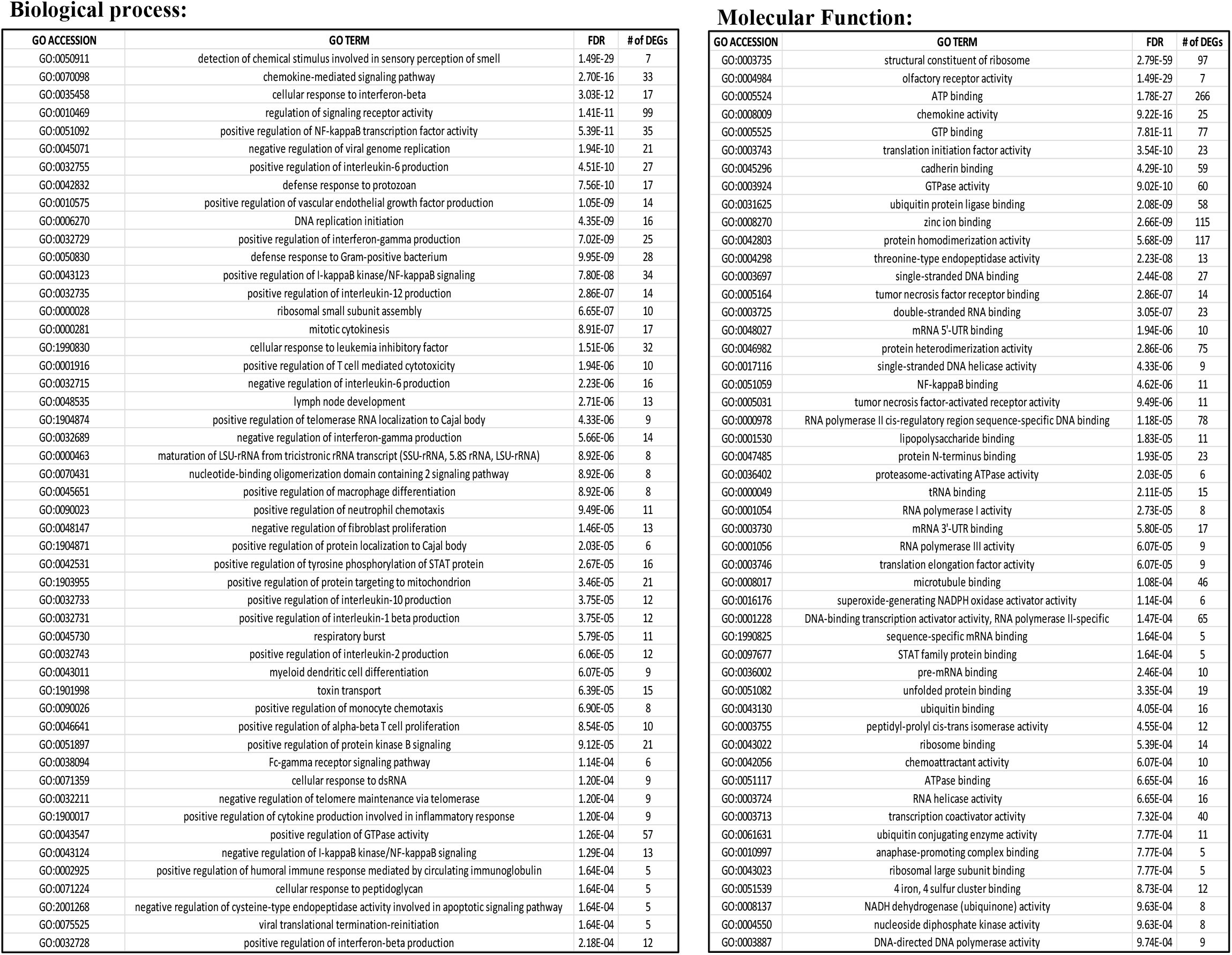
Significant upregulated GO terms. Gene Ontology analysis of the differentially expressed genes in lungs of LDR mice compared to naïve mice, ranked by false discovery rate (FDR) after eliminating redundant terms. Fisher’s exact test was carried out to estimate the association between differentially expressed genes with adjusted *p* value < 0.05 and specific GO categories using the OmicsBox software. Presented are the top 50 GO terms of biological process and molecular function terms, with their FDR value and the number DEGs.

**Fig. S2:**
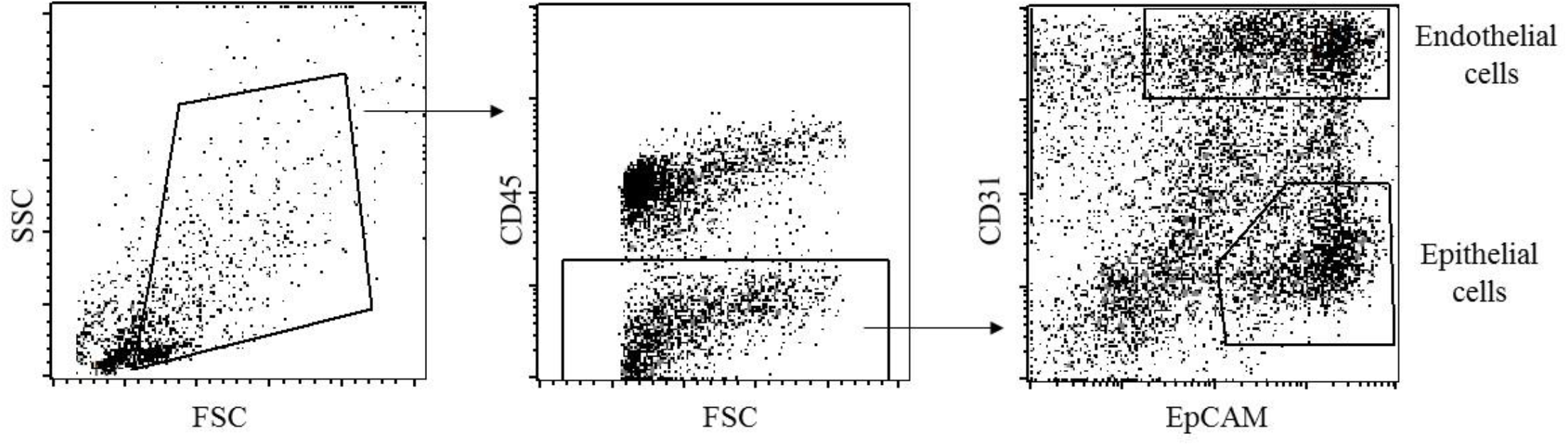
A hierarchical gating strategy to identify lung epithelial and endothelial cells. Representative FACS plot that identifies lung epithelial cells (CD45-CD31-EpCAM+) and endothelial cells (CD45-CD31+).

## References

1. F. Zhou, T. Yu, R. Du, G. Fan, Y. Z. Liu, J. Xiang, Y. Wang, B. Song, X. Gu, L. Guan, Y. Wei, H. Li, X. Wu, J. Xu, S. Tu, Y. Zhang, H. Chen, B. Cao. Clinical course and risk factors for mortality of adult inpatients with COVID-19 in Wuhan, China: a retrospective cohort study. Lancet 395, 1054–162 (2020).

2. G. Grasseli, A. Zangrillo, A. Zanella, M. Antonelli, L. Cabrini, A. Castelli, D. Cereda, A. Coluccello, G. Foti, R. Fumagalli, G. Iotti, N. Latronico, L. Lorini, S. Merler, G. Natalini, Alessandra Piatti, Marco Vito Ranieri, A. Mara Scandroglio, E. Storti, M. Cecconi, A. Pesenti. Baseline characteristics and outcomes of 1591 patients infected with SARS-CoV-2 admitted to ICUs of the Lombardy region, Italy. JAMA 323, 1574–1581 (2020).

3. Y.-I. Kim, S.G. Kim, S.M. Kim, E.H Kim, S.J. Park, K.M. Yu, J.H. Chang, E.J. Kim, S. Lee, M.A.B. Casel, J. Um, M.S. Song, H.W. Jeong, V.D. Lai, Y. Kim, B.S. Chin, J.S. Park, K.H. Chung, S.S. Foo, H. Poo, I.P. Mo, O.J. Lee, R.J. Webby, J.U. Jung, Y.K. Choi. Infection and rapid transmission of SARS-CoV-2 in ferrets. Cell Host Microbe. 27, 704–709 (2020).

4. Y. Yahalom-Ronen, H. Tamir, S. Melamed, B. Politi, O. Shifman, H. Achdout, E. B. Vitner, O. Israeli, E. Milrot, D. Stein, I. Cohen Gihon, S. Lazar, H. Gutman, I. Glinert, L. Cherry, Y. Vagima, S. Lazar, S. Weiss, A. Ben-Shmuel, R. Avraham, R. Puni, E. Lupu, E. Bar David, A. Sittner, N. Erez, R. Zichel, E. Mamroud, O. Mazor, H. Levy, O. Laskar, S. Yitzhaki, S. C. Shapira, A. Zvi, A. Beth-Din, N. Paran, T. Israely. A single dose of recombinant VSV-ΔG-spike vaccine provides protection against SARS-CoV-2 challenge. BioRxiv (2020). https://doi.org/10.1101/2020.06.18/160655.

5. B. Rockx, T. Kuiken, S. Herfst, T. Bestebroer, M.M. Lamers, B.B. Oude Munnink, D. de Meulder, G. van Amerongen, J. van den Brand, N.M.A. Okba, D. Schipper, P. van Run, L. Leijten, R. Sikkema, E. Verschoor, B. Verstrepen, W. Bogers, J. Langermans, C. Drosten, M. Fentener van Vlissingen, R. Fouchier, R. de Swart, M. Koopmans, B.L. Haagmans. Comparative pathogenesis of COVID-19, MERS, and SARS in a nonhuman primate model. Science 3689:1012–1015 (2020).

6. L. Bao, W. Deng, B. Huang, H. Gao, J. Liu, L. Ren, Q. Wei, P. Yu, Y. Xu, F. Qi, Y. Qu, F. Li, Q. Lv, W. Wang, J. Xue, S. Gong, M. Liu, G. Wang, S. Wang, Z. Song, L. Zhao, P. Liu, L. Zhao, F. Ye, H. Wang, W. Zhou, N. Zhu, W. Zhen, H. Yu, X. Zhang, L. Guo, L. Chen, C. Wang, Y. Wang, X. Wang, Y. Xiao, Q. Sun, H. Liu, F. Zhu, C. Ma, L. Yan, M. Yang, J. Han, W. Xu, W. Tan, X. Peng, Q. Jin, G. Wu, C. Qin. The pathogenicity of SARS-CoV-2 in hACE2 transgenic mice. Nature 583, 830–840 (2020).

7. A.O. Hassan, J.B. Case, E.S. Winkler, L.B. Thackray, N.M. Kafai, A.L. Bailey, B.T. McCune, J.M. Fox, R.E. Chen, W.B. Alsoussi, J.S. Turner, A.J. Schmitz, T. Lei, S. Shrihari, S.P. Keeler, D.H. Fremont, S. Greco, P.B. McCray Jr, S. Perlman, M.J. Holtzman, A.H. Ellebedy, M.S. Diamond. A SARS-CoV-2 infection model in mice demonstrates protection by neutralizing antibodies. Cell 182, 1–10 (2020).

8. E.S. Winkler, A.L. Bailey, N.M. Kafai, S. Nair, B.T. McCune, J. Yu, J.M. Fox, R.E. Chen, J.T. Earnest, S.P. Keeler, J.H. Ritter, L.I. Kang, S. Dort, A. Robichaud, R. Head, M.J. Holtzman, M.S. Diamond. SARS-CoV-2 infection of human ACE2-transgenic mice causes severe lung inflammation and impaired function. Nature Imm. (2020). doi:10.1038/s41590-020-0778-2

9. G. Matute-Bello, C.W. Frevert, T.R. Martin. Animal models of acute lung injury. Am. J. Physio. Lung Cell Mol. Physiol. 295, L379–L399 (2008).

10. S. Katalan, R. Falach, A. Rosner, M. Goldvaser, T. Brosh-Nissimov, A. Dvir, A. Mizrachi, O. Goren, B. Cohen, Y. Gal, A. Sapoznikov, S. Ehrlich, T. Sabo, C. Kronman. A novel swine model of ricin-induced respiratory distress syndrome. Dis. Models Mech. 10, 173–183 (2017).

11. J. Radbel, D.L. Laskin, J.D. Laskin, H.M. Kipen. Disease-modifying treatment of chemical threat agent-induced acute lung injury. Ann. N.Y. Acad. Sci. (2020). doi:10.1111/nyas.14438

12. P. Zhou, X.L. Yang, X.G. Wang, B. Hu, L. Zhang, W. Zhang, H.R. Si, Y. Zhu, B. Li, C.L. Huang, H.D. Chen, J. Chen, Y. Luo, H. Guo, R.D. Jiang, M.Q. Liu, Y. Chen, X.R. Shen, X. Wang, X.S. Zheng, K. Zhao, Q.J. Chen, F. Deng, L.L. Liu, B. Yan, F.X. Zhan, Y.Y. Wang, G.F. Xiao, Z.L. Shi. A pneumonia outbreak associated with a new coronavirus of probable bat origin. Nature 579, 270–275 (2020).

13. Y. Vagima, E. Grauer, B. Politi, S. Maimon, E. Yitzhak, S. Melamed, H. Achdout, D. Gur, M. Aftalion, A. Shemesh, A. Hasson, S. Yitzhaki, S.C. Shapira, E. Mamroud. Group activity of mice in communal home cage used as an indicator of disease progression and rate of recovery: effects of LPS and influenza virus. Life Sci. 258, 118214 (1-8) (2020).

14. R. Falach, A. Sapoznikov, Y. Evgy, M. Aftalion, A. Makovitzki, A. Agami, A. Mimran, E. Lerer, A. Ben David, R. Zichel, S. Katalan, A. Rosner, T. Sabo, C. Kronman, Y. Gal. Post-exposure ant-ricin treatment protects swine against lethal systemic and pulmonary exposures. Toxins 12, 354 (1-11) (2020).

15. Y. Gal, O. Mazor, R. Alcalay, N. Seliger, M. Aftalion, A. Sapoznikov, R. Falach, C. Kronman, T. Sabo. Antibody/doxycycline combined therapy for pulmonary ricinosis: attenuation of inflammation improves survival of ricin-intoxicated mice. Toxicol. Rep. 1, 496–504 (2014).

16. A. Mechaly, U. Elia, R. Alcalay, H. Cohen, E. Epstein, O. Cohen, O. Mazor. Inhibition of Francisella tularensis macrophage-uptake by a novel anti-LPS scFv antibody fragment. Sci. Rep. 9, 11418–11426 (2019).

17. M. Hoffmann, H. Kleine-Weber, S. Schroeder, N. Krüger, T. Herrler, S. Erichsen, T.S. Schiergens, G. Herrler, N.H. Wu, A. Nitsche, M.A. Müller, C. Drosten, S. Pöhlmann. SARS-CoV-2 cell entry depends on ACE2 and TMPRSS2 and is blocked by a clinically proven protease inhibitor. Cell 181, 271–280 (2020).

18. T. Noy-Porat, E. Makdasi, R. Alcalay, A. Mechaly, Y. Levy, A. Bercovich-Kinori, A. Zauberman, H. Tamir, Y. Yahalom-Ronen, M. Israeli, E. Epstein, H. Achdout, S. Melamed, T. Chitlaru, S. Weiss, E. Peretz, O. Rosen, N. Paran, S. Yitzhaki, S.C. Shapira, T. Israely, nO. Mazor, R. Rosenfeld. A panel of human neutralizing mAbs targeting SARS-CoV-2 spike at multiple epitopes. Nature Comm. (2020). https://doi.org/10.1038/s41467-020-18159-4.

19. T. Noy-Porat, O. Cohen, S. Ehrlich, E. Epstein, R. Alcalay, O. Mazor. Acetylcholinesterase-Fc Fusion protein (AChE-Fc): a novel potential organophosphate bioscavenger with extended plasma half-life. Bioconj. Chem. 26, 1753–1758 (2015).

20. K.S Corbett, D. Edwards, S.R. Leist, O.M. Abiona, S. Boyoglu-Barnum, R.A. Gillespie, S. Himansu, A. Schäfer, C.T. Ziwawo, A.T. DiPiazza, K.H. Dinnon, S.M. Elbashir, C.A. Shaw, A. Woods, E.J. Fritch, D.R. Martinez, K.W. Bock, M. Minai, B.M. Nagata, G.B. Hutchinson, K. Bahl, D. Garcia-Dominguez, L. Ma, I. Renzi, W.P. Kong, S.D. Schmidt, L. Wang, Y. Zhang, L.J. Stevens, E. Phung, L.A. Chang, R.J. Loomis, N.E. Altaras, E. Narayanan, M. Metkar, V. Presnyak, C. Liu, M.K. Louder, W. Shi, K. Leung, E.S. Yang, West, K.L. Gully, N. Wang, D. Wrapp, N.A. Doria-Rose, G. Stewart-Jones, H. Bennett, M.C. Nason, T.J. Ruckwardt, J.S. McLellan, M.R. Denison, J.D. Chappell, I.N. Moore, K.M. Morabito, J.R. Mascola, R.S. Baric, A. Carfi, B.S. Graham. SARS-CoV-2 mRNA vaccine design enabled by prototype pathogen preparedness. Nature (2020). doi:10.1038/s41586-020-2622-0

21. H. Gu, Q. Chen, G. Yang, L. He, H. Fan, Y.Q. Deng, Y. Wang, Y. Teng, Z. Zhao, Y. Cui, Y. Li, X.F. Li, J. Li, N.N. Zhang, X. Yang, S. Chen, Y. Guo, G. Zhao, X. Wang, D.Y. Luo, H. Wang, X. Yang, Y. Li, G. Han, Y. He, X. Zhou, S. Geng, X. Sheng, S. Jiang, S. Sun, C.F. Qin, Y. Zhou. Adaptation of SARS-CoV-2 in Balb/C mice for testing vaccine efficacy. Science (2020) 10.11.26/science/abc4730 (2020).

22. V. G. Puelles, M. Lütgehetmann, M.T. Lindenmeyer, J.P. Sperhake, M.N. Wong, L. Allweiss, S. Chilla, A. Heinemann, N. Wanner, S. Liu, F. Braun, S. Lu, S. Pfefferle, A.S. Schröder, C. Edler, O. Gross, M. Glatzel, D. Wichmann, T. Wiech, S. Kluge, K. Pueschel, M. Aepfelbacher, T.B. Huber. Multiorgan and renal tropism of SARS-CoV-2. N. Engl. J. Med. 383, 590–592 (2020).

23. Y.-Y. Zheng, Y.T. Ma, J.Y. Zhang, X. Xie. COVID-19 and the cardiovascular system. Nature Rev. Cardiol. 17, 259–260 (2020).

24. M. Mateo, A. Generous, P.L., Sinn, R. Cattaneo. Connections matter-how viruses use cell-cell adhesion components. J. Cell Sci. 128, 431–439 (2015).

25. A. Sapoznikov, Y. Gal, R. Falach, I. Sagi, S. Ehrlich, E. Lerer, A. Makovitzki, A. Aloshin, C. Kronman, T. Sabo. Early disruption of the alveolar-capillary barrier in a ricin-induced ARDS mouse model: neutrophil-dependent and -independent impairment of junction proteins. Am. J. Physiol. Cell Mol. Physiol. 316, L255–L268 (2019).

26. H. Ohta, S. Chiba, M. Ebina, M. Furuse, T. Nukiwa. Altered expression of tight junction molecules in alveolar septa in lung injury and fibrosis. Am. J. Physiol. Cell Mol. Physiol. 302, L193–L205 (2012).

27. S.M. Evans, D.I. Blyth, T. Wong, S. Sanjar, M.R. West. Decreased distribution of lung epithelial junction proteins after intratracheal antigen or lipopolysaccharide challenge: correlation with neutrophil influx and levels of BALF sE-cadherin. Am. J. Respir. Cell Mol. Biol. 27, 446–454 (2002).

28. R. Falach, A. Sapoznikov, Y. Gal, O. Israeli, M. Leitner, N. Seliger, S. Ehrlich, C. Kronman, T. Sabo. Quantitative profiling of the in vivo enzymatic activity of ricin reveals disparate depurination of different pulmonary cell types. Toxicol Lett. 258, 11–19 (2016).

29. E. Keyaerts, L. Vijgen, P. Maes, J. Neyts, M. Van Ranst. Growth kinetics of SARS-coronavirus in Vero E6 cells. Biochem. Biophys. Res. Comm. 329, 1147–1151 (2005).

